# Preliminary Evidence of the role of estrogen and tamoxifen-induced regulation of complement proteins in rat hippocampus

**DOI:** 10.1101/2020.01.30.927392

**Authors:** Pavan Kumar, Pushpa Dhar

## Abstract

Effects of Estrogen (E2) is widespread in the human body; still, an unresolved paradox. Neurodegeneration and neuroinflammation are inherently associated with age progression, debilitating by hormone deprivation, especially in female. Senescent cells accumulate with age and promote tissue deterioration in the body system. Neurodegenerative diseases drive a healthy life towards to morbidity and feebleness; despite the different etiology, uncontrolled inflammation is one of the significant causals factors. We here used post-menopausal model (ovariectomized female rat), E2 replenishment therapy reduces the expression of inflammatory mediators, such as complement proteins (C3, C1q, and C3aR) in these animals.

E2 therapy could limit the ovariectomy-induced increase of inflammatory events in brain regions such as the hippocampus. Also, the duration of hormone deprivation could be a determinant for the intensity of the anti-inflammatory actions of estrogen. On the whole, considerable evidence, including that from the present study supports the view that complement biosynthesis, which plays a significant role in phagocytosis of cellular debris and synaptic pruning of postnatal neural circuits goes uncontrolled and could be the inducing factor for enhanced neurodegeneration following hormone deprivation.

## 1. Introduction

Estrogen, a cholesterol derivative, belongs to the steroid hormone family. It is majorly synthesized in the ovaries and to a lesser extent in the testes and placenta. Apart from its profound influence on sexual differentiation and function, estrogen plays a definitive role in modulating the functioning of various other systems such as cardiovascular, skeletal, nervous, etc. In the brain, estrogen acts not only on the hypothalamo-hypophyseal axis but also in areas like the hippocampus, frontal cortex, cerebellum, etc. thereby influencing the brain functions that are not related to reproductive activity directly. In these brain areas, estrogen shows modulatory influence on associated functions like memory cognition, postural stability, skilled motor activity, etc. Estrogen has also been reported as a potent neuroprotectant in various experimental models (Dubal et al., 1998; Green and Simpkins, 2000; Gridley et al., 1997; McEwen, 2002; O’Neal et al., 1996). The biological effects of estrogen in the CNS range from induction of neurogenesis (Tanapat et al., 2005; Tanapat et al., 1999) to conferring neuroprotective effects (Garcia-Segura et al., 2001). Estrogen acts through specific nuclear receptors, ERα and ERβ (Kuiper et al., 1996; Mosselman et al., 1996) to influence the healthy development and differentiation of the brain, as well as the production and maintenance of sexually dimorphic behaviour throughout the adult life (Breedlove, 1992; Jones et al., 1988). Activated ERs act as transcription factors that bind to specific DNA sequences of target genes, the Estrogen Response Elements (EREs) (Gruber et al., 2004).

Acute and chronic inflammation in different brain areas is believed to influence various functions such as learning, memory, etc., besides having an impact on age-related neurodegenerative conditions. The first act of the innate immune system is to the recruitment of immune cells at the site of damage by releasing chemoattractant, followed by clearing the dead cells and foreign invaders. The complement system, an essential component of the innate immune response, is activated via three pathways: classical, lectin-dependent, and alternative pathways. Levi-Strauss and Mallat (1987) were the first to report the release of complement proteins by brain cells. Increase in local complement protein biosynthesis and their uncontrolled activation in the CNS may augment the neurodegenerative pathology (Levi-Strauss and Mallat, 1987; Ramaglia et al., 2008). Decreased estrogen levels during menopause are reported to increase the inflammatory risk (Abu-Taha et al., 2009; Barrientos et al., 2019). There are reports of modulation of the microglial activation and other inflammatory molecules, including complement component in the rat brain following estrogen replenishment therapy (Sarvari et al., 2011; Sarvari et al., 2014). The proposed mechanisms of neuroprotective effects augmented by estrogen might include its anti-inflammatory responses (innate immune response), which in turn could influence the neurodegenerative pathology.

Tamoxifen is a synthetic non-steroidal compound and has tissue-specific effects distinct from those of estradiol, acting as estrogen receptor agonists in some tissues and as antagonists in others (Riggs and Hartmann, 2003; Smith and O’Malley, 2004). Because of its selective nature, it’s also known as a selective estrogen modulator (SERM). Like estrogen, tamoxifen binds to the classical receptors (ERα and ERβ) and elicits rapid membrane and cytoplasmic actions by membrane initiated steroid signaling. Tamoxifen exerts its protective effect by triggering phosphorylation of MAPK/ERK CREB and PI3K/Akt/mTOR signaling pathways and reducing inflammation, lipid peroxidation, ROS formation, Aβ plaques, and apoptotic proteins. Kimelberg and colleagues (2000) were the first to report a neuroprotective action of tamoxifen in animal (male) models of stroke, an effect later extended to female animals by other investigators (Mehta et al., 2003).

Hence, the present study aimed at determining the immunoregulatory and neuroprotective role of estrogen and Tamoxifen (SERM) supplementation on the hippocampus of ovariectomized (hormone depleted) rats.

## 2. Materials and Methods

### 2.1. Animals

Female Wistar rats (Rattus norvegicus), 4-5 months old with 3-4 consecutive regular estrus cycles, were included in the study (n=60). The animals were procured from the Institutional Animal Facility with the approval of the Ethics Committee (IAEC) and housed in the same facility under controlled environmental conditions (12:12 light-dark cycle) with and libitum access to food and water. The instructions for care of the animals as laid down by the IAEC were strictly followed.

### 2.2. Vaginal smear cytology

For determining the status of the estrous cycle of the animals, vaginal smears were taken daily (before noon) following the standard procedure (Markowska, 1999). After staining with Wright-Giemsa method (Sigma Aldrich, USA), the smears were observed under the bright field microscope and staged for the phases of the estrous cycle.

### 2.3. Ovariectomy (OVX)

During the estrous phase of the estrous cycle, bilateral ovariectomy was performed under all aseptic conditions via a dorsal incision in the flank area (Zhou et al., 2002). After a recovery time of two weeks following surgical trauma, the vaginal smear studies were undertaken, and complete absence of mature epithelial cells in the smears confirmed their passing into a persistent Diestrus phase and hence the successful ovariectomy. This was followed by the designed treatment plans for 30 days.

### 2.4. Animal groups

The animals were divided into five groups (Grps) (n=12/grp). Ovary intact animals (OVI) served as normal controls (Grp I), and ovariectomized (OVX) animals served as OVX controls (Grp II). The animals (OVX) in Groups III to V received subcutaneous (s.c.) injection of vehicle (0.1 ml Sesame oil), 17β-estradiol (E2, 0.1 mg/kg) and TAM (0.05 mg/kg) respectively for 30 days. The dose selection was according to previously published reports (Mehra et al., 2005; Sharma and Mehra, 2008; Sharma et al., 2007).

### 2.5. Morris water maze test

Memory is a collectively a set of diverse cognitive capacities by which humans and animals retain information and recollect past experiences. Spatial memory in rodents involves the ability to map the environment in three-dimensional space. The Morris water maze has proven to be a robust and reliable test that is strongly correlated with hippocampal synaptic plasticity and hippocampal-dependent spatial learning and memory in rats (Morris, 1984). A circular pool (210 cm in diameter; 51cm deep) was filled with water 25^0^-27^0^C upto the level of 21 cm from bottom. A removable white plexiglass platform (11cm diameter) keept in the Southwest (SW) quadrant of the pool was used as escape platform from the water. Animals were tested for four consecutive days (4 trials/day). During each acquisition trial the animal was placed into the water (60 sec) and allowed to locate platform, parameters such as escape latency and distance travelled were determined by NI-IMAQ software (Coulbourne Instruments, USA). To assess reference memory probe trials were given after 24 hrs of the last acquisition trial. If, the animal failed to find the escape platform during maximum allocated time, the animal was guided to the platform by the experimenter with the help of a ruler. The animal was allowed to remain on the platform for 15 seconds.

### 2.6. Immunohistochemical (IHC) studies

Brain tissue obtained from perfusion fixed animals (n=6) was processed for cryo-sectioning (30μm) followed by IHC and IF process. The protocols of previously published reports for IHC (Mehra et al., 2005; Sharma and Mehra, 2008; Sharma et al., 2007) were followed. Complement C3 specific mouse monoclonal (Sc-28294, RRID:AB_627277), Complement C1q specific rabbit polyclonal (Sc-25856, RRID:AB_2228238), C3aR (Sc-20138, RRID:AB_2066739), Iba1 (ab5076, RRID:AB_2224402) and SYP (Sc-9116, RRID:AB_2199007) were used as primary antibodies (1:200). Anti-polyvalent HRP-Streptavidin Kit (Thermo Scientific) with DAB was used for the visualization of antigenic sites.

### 2.7. Western blot analysis and qRTPCR

Fresh brain tissue (n=6) was used for proteomics and gene expression (qRT-PCR) analysis. The hippocampus was separated in RNAse free condition and left in RNAlater at room temperature (RT) and later stored at -80^0^C. Protein and RNA extraction was done using Qiagen AllPrep DNA/RNA/Protein isolation kit (Qiagen Inc, USA). RNA purity was confirmed with 260/280 OD ratio using Nanodrop reader (Thermo Scientific, USA) and total tissue protein was estimated with the Bradford method (Bradford 1976). Supernatants of all the samples were used for Western blot analysis (Mehra et al., 2005; Sharma and Mehra, 2008; Sharma et al., 2007). Primary antibodies (C3, C3aR, C1q and loading control beta actin ab75186 RRID:AB_1280759) and IgG-HRP conjugate secondary antibodies (Sc-2005, RRID:AB_631736 and Sc-2030 RRID:AB_631747, and SC-2033 RRID:AB_631729; Santa Cruz, USA) were used at a dilution of 1:500 & 1:1000 respectively. Densitometric analysis of protein bands was done by Imagestudio Lite software (LI-COR Biosciences, USA) For qRT-PCR, first-strand cDNA synthesis from total RNA templates present in samples (hippocampus) was done by RevertAid First Strand cDNA Synthesis Kit (Thermo Scientific). Negative (no RT and no template) and positive (reference gene β-actin and GAPDH) controls were included in the experiment. Gene expression was determined using Maxima SYBR green qRT-PCR kit (Thermo Scientific, USA) and gene-specific primers. In brief, a qRT-PCR assay was designed for genes of interest (C3 and C1q) and reference genes (β Actin and GAPDH). Real-time qPCR was performed in a 96 -well plate format using BIOER Linegene thermal cycler (BIOER, Hong Kong). Ct values obtained were normalized with the reference gene, and relative fold expression was calculated using the Livak method (ΔΔCT). Primer sequences were as follows: for C3, forward 5-CTGTACGGCATAGGGATATCACG-3, reverse 5-ATGCTGGCCTGACCTTCAAGA-3 (Misumi et al., 1990); for C1q, forward 5-ACCCAGACTTCCGCTTTCTG-3, reverse 5-CTTCAGGGGCCTCCTGTGTA-3 (self-constricted); for β actin, forward 5-GCGCTCGTCGTCGACAACGG-3, reverse 5-GTGTGGTGCCAAATCTTCTCC-3 (Kit provided); for GAPDH, forward 5-CAAGGTCATCCATGACAACTTTG-3, reverse, 5-GTCCACCACCCTGTTGCTGTAG-3 (Lin et al., 2013); for C3a forward 5-GACCTACACTCAGGGC-3, reverse 5-ATGACG GACGGGATAAG-3 (Benard et al., 2004); for forward SYP 5-TCAGGACTCAACACCTCAGTGG-3 reverse 5-AACACGAACCATAAGTTGCCAA-3 (Chen et al., 2001).

### 2.8. Serum estradiol assay

Twenty-four hours after the last injection, blood samples were obtained from the tail vein, and serum was stored at – 20 °C for subsequent estradiol estimations (Estradiol 17β Serum/Plasma EIA kit; Assay designs, USA, Cat. No. 900-174).

At the end of the experimental period, the animals were deeply anesthetized and either subjected to perfusion fixation or cervical dislocation.

### 2.9 TUNEL (Terminal deoxynucleotidyl transferase dUTP nick end labelling) assay

TUNEL assay (Neuro TACS in Situ Apoptosis Detection Kit, Trevigen) was carried out on 10μm thick cryosections, and apoptotic cells were identified by the presence of reddish-brown nuclear precipitate in DAB reaction. The sections were counterstained with CV for the demarcation of the cell bodies. The TUNEL positive neurons were counted in sampling fields (280 μm × 380 μm) and the data from individual animals/group were pooled to calculate the percentage of apoptotic cells (Sharma et al., 2007).

### 2.10. Quantitative analysis

For the quantitative analysis, Integrated Optical Density (IOD) was measured. In brief, five (coronal) sections/animal/group were analysed with the help of Image Pro-Plus software (v 6.2, Media Cybernetics, USA).

### 2.11. Statistical analysis

Statistical analysis was done by using Graphpad prim 7. One-way analysis of variance (ANOVA) with post hoc multiple comparisons using the students Neuman-Keuls test and Two-way ANOVA with Bonferroni post-tests were carried out. Data from individual animals/group were pooled together, and the results were expressed as mean ± SEM.xs

## 3. Results

### 3.1. Memory impairment in OVX rat

All animals were given training on the first day to get acclimatized to the water maze task. Later, 4 trial per day for 4 consecutive days were carried out for the spatial acquisition of memory in the experimental and the control rats. During the acquisition phase, the distance traveled (cm) and time taken to reach the platform (sec) were recorded as measure of performance task in the water maze. The distance traveled by animals decreased with continued training [F (3, 220) = 745.45; P<0.0001], thus reflecting the ongoing process of learning to reach the platform by travelling shorter distances. Following ovariectomy, the distance traveled by animals to reach the platform on day 1, and day 4, was significantly increased as compared to ovary intact controls. However, the distance traveled by OVX animals receiving E2 and TAM treated animals (compared to OVX controls) was significantly lesser than ovariectomized controls, thus reflecting a better performance on spatial memory by former groups. Vehicle treatment to OVX rats did not show any significant effect on distance traveled and hence, the performance was more or less similar to that of the ovariectomized controls (Fig 1A).

**Fig. 1.**
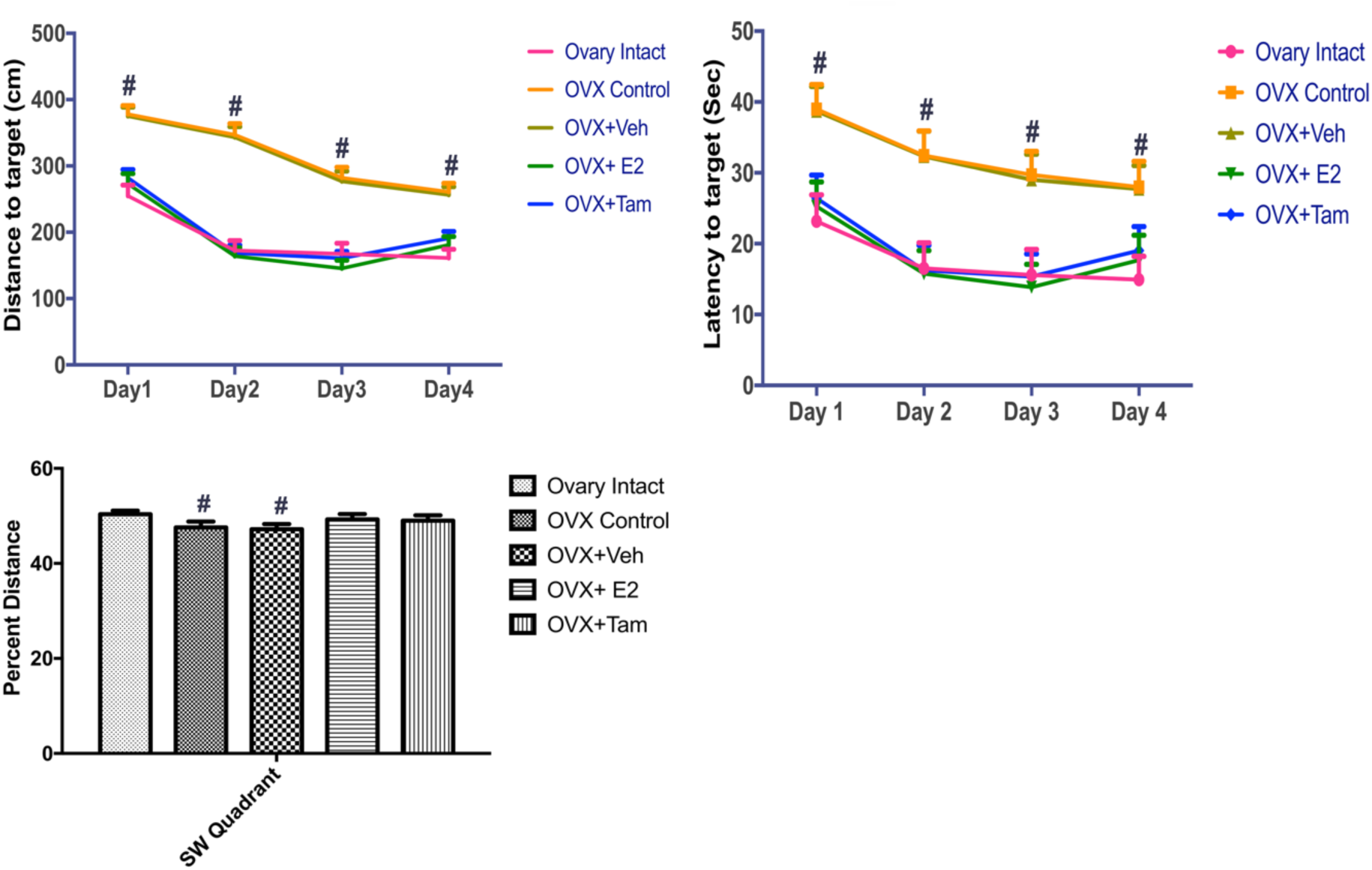
Memory impairment is increased in ovariectomized rat. Line graph summarising the effects of E2 depletion (OVX) and treatment with E2 and SERM (TAM) on hippocampal spatial memory of rats assessed as distance swam (A), escape latency and percentage distance traveled in the target zone (C) were recorded. Values are expressed as Mean±SEM; n=12, #p<0.001 in comparison to OVX controls. Two-way ANOVA with Bonferroni posttest.

The escape latency also decreased across the days [F (3, 220) = 102.8; P<0.0001]. Following ovariectomy, the animals took a longer time to locate and reach the platform, hence the escape latency was significantly increased both on day 1and day 4 when compared to ovary intact controls. However, hormone and SERM (E2 and TAM) treatments significantly decreased the escape latency in OVX+E2 and OVX+TAM compared to OVX controls. Vehicle treatment had no significant effect on escape latency and was comparable to OVX controls (Fig. 1B). The probe (transfer) trial, to assess reference memory [F (4, 55) = 16.82; P<0.0001], 24 h after the last acquisition day. Observed data revealed that OVX controls covered significantly less distance in the SW quadrant (original platform position) than that covered by ovary intact controls in the same quadrant (Fig. 2C). E2 treated and TAM treated animals traveled more distance in SW quadrant. However, distance traveled in a SW quadrant by vehicle-treated rats was similar to OVX rats. These observations are suggestive of spatial reference memory induced by hormone depletion. However, hormone or SERM therapy resulted in an improvement of reference reflected by their preference of spending more time in platform quadrant (Fig. 3).

**Fig. 2.**
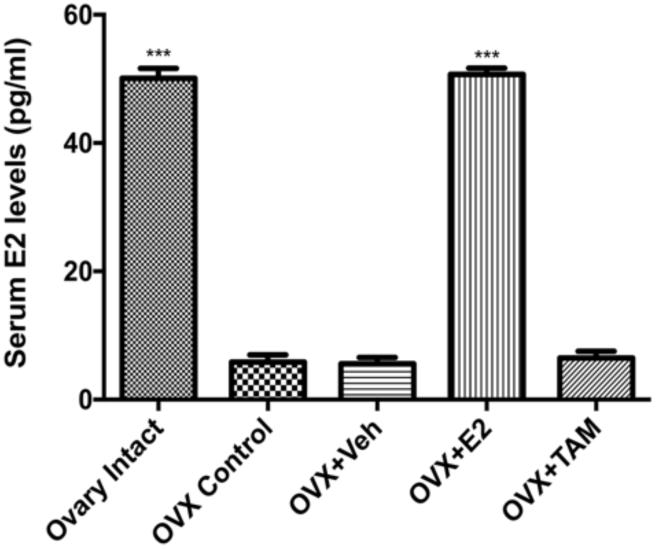
Bardigram showing serum estradiol (E2) levels (pg/ml) (mean ±SEM). Serum estradiol level in OVI and E2 treated OVX rats were significantly (***p<0.001) higher, the levels being comparable amongst OVX control, OVX+Veh and OVX+TAM treated groups.

**Fig. 3.**
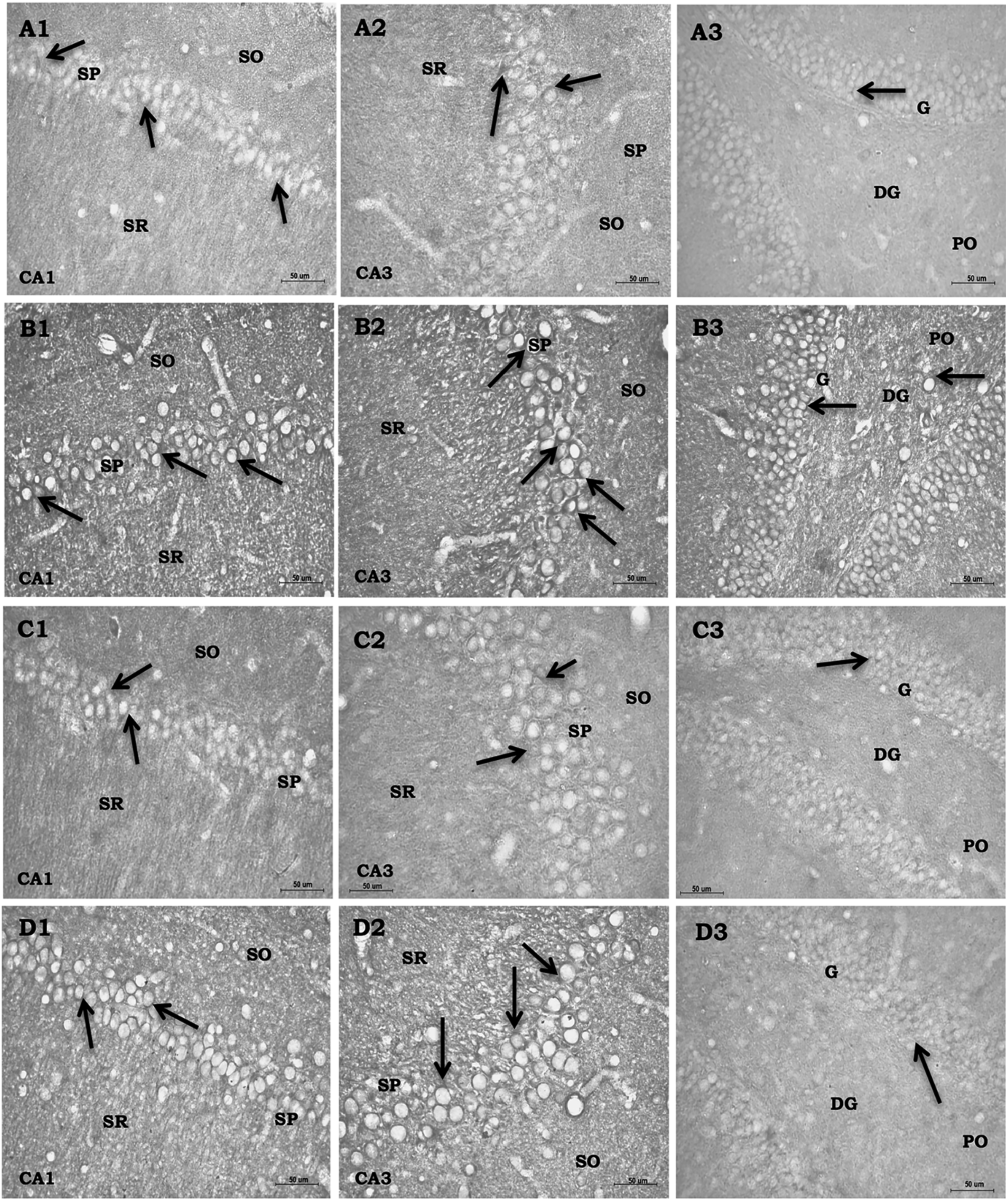
High-power photomicrographs of coronal sections of rat hippocampus (CA1, CA3 & DG) of OVI (A1-A3), OVX (B1-B3), OVX +E2 (C1-C3) and OVX +TAM (D1-D3) showing distribution of complement C3 +ve neurons. Ovariectomy led to significant upregulation of C3 irty in perikaryal membrane (B1-B3). Whereas E2 or TAM therapy to OVX led to significant downregulation of C3 in all the subfields of hippocampus (C1-C3; C1-D3).

### 3.2. Serum estradiol levels

Serum E2 (pg/ml) level [F (4, 25) = 547.8; P<0.0001; η2= 0.9887] was significantly lower in OVX (5.85±1.12; p< 0.0001) and OVX+Veh (5.58±0.97; p< 0.0001) animals as compared to ovary intact controls (42.41±0.89). E2 administration to OVX animals led to a significant increase in serum E2 levels (49.02±0.51; p< 0.0001), being comparable to those of ovary intact controls (OVI). TAM treatment had no significant effect on serum E2 levels (6.48±1.03; p< 0.0001) (Fig. 2).

### 3.3. Immunohistochemical expression of complement proteins (C3, C3aR, and C1q)

#### 3.3.1. C3 expression

Expression of complement protein C3 was exclusively localized in the perikaryal membrane and was observed in all the subfields (CA1, CA3, and DG) of the hippocampus in OVI animals with a significant proportion of C3 +ve cells in the SP of CA1 and CA3 (Fig. 3A1 & A2) and granular (G) and polymorphic (PO) layers of DG (Fig. 3A3). C3 immunoreactivity was significantly upregulated in all the regions following ovariectomy (Fig 3B1, B2, and B3). E2 and TAM therapy in OVX animals resulted in downregulation of C3 expression in the SP (CA1 and CA3) (Fig. 3C1, C2; D1, D2), the intensity being comparable to ovary intact-like state (Fig. 3A1, A2). Relatively low expression of C3 was also observed in granular and polymorphic cell layers of DG in E2 and TAM supplemented groups (Fig. 3C3, D3).

#### 3.3.2. C3aR expression

Expression of C3aR was localized in the perikarya as well as processes (dendrites and axons) of hippocampal neurons. In OVI animals, C3aR+ve neurons were mainly localized in SP and SO of CA1 & CA3 (Fig. 4A1, A2) whereas C3aR irty in DG (G & PO layers) of OVI animals (Fig. 4A3) was quite low. A robust increase in C3aR expression was seen in SP and SO (CA1 & CA3) (Fig. 4B1, B2) and G & PO layers of DG (Fig. 4B3) of OVX animals. E2 and TAM therapy to OVX animals led to a significant downregulation of C3aR irty in CA1 and CA3 (Fig. 4C1-C2 & D1, D2). Whereas, the intense expression of C3aR was evident in cell layers of DG in TAM treated animals (Fig. 4D3).

**Fig. 4.**
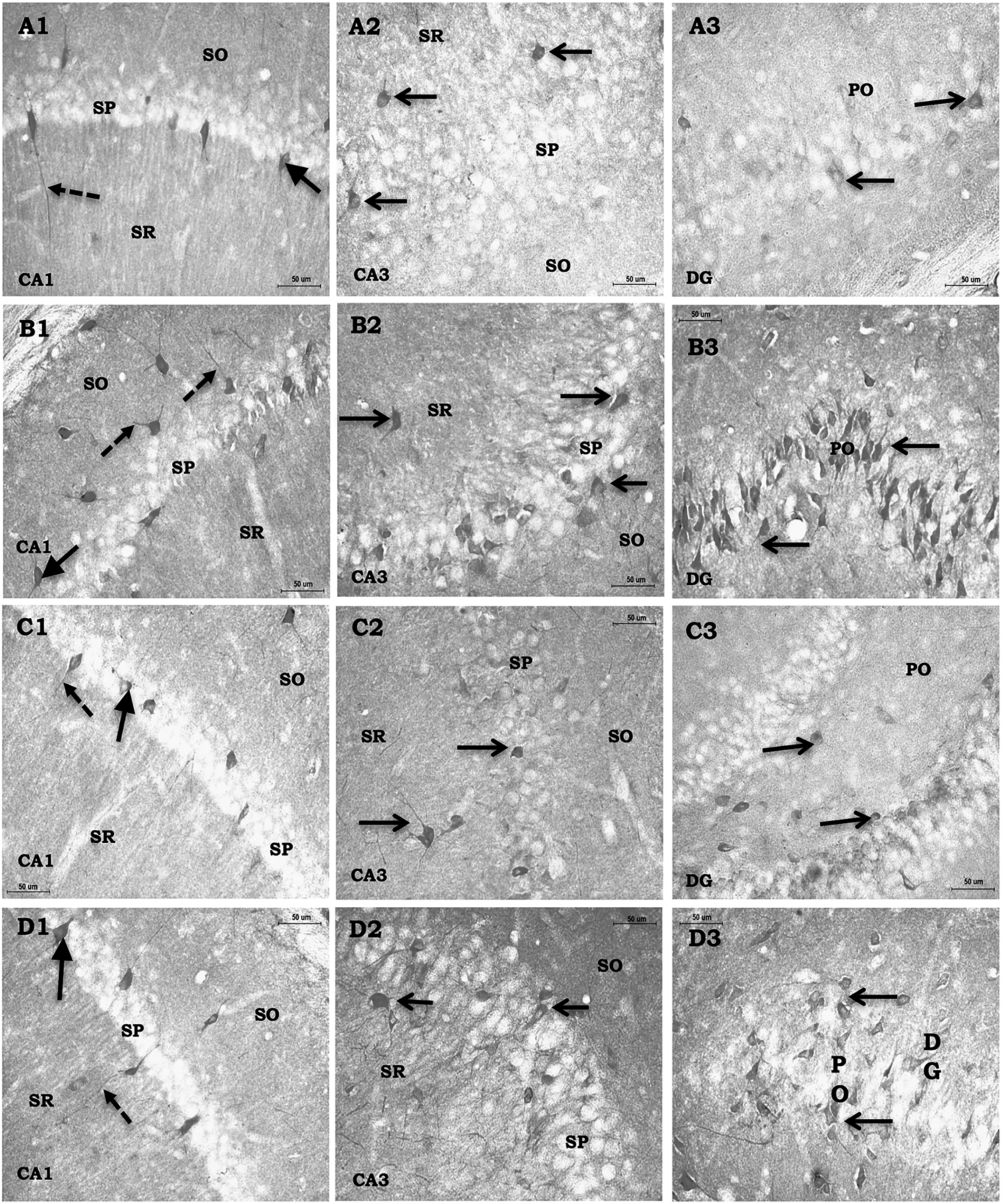
High-power photomicrographs of coronal sections of rat hippocampus of OVI (A1-A3), OVX (B1-B3), OVX +E2 (C1-C3) and OVX + TAM (D1-D3) showing distribution of complement C3aR +ve neurons. Note: -C3aR irty was observed in perikaryal membrane (red aroow), dendritic and axonal process (green aroow) of most of the neurons in SP, SO & PO (black arrows). Significant upregulation of C3aR irty was observed following ovariectomy (B). Whereas E2 (C) or TAM (D) therapy to OVX rats leads to significant downregulation of C3aR, being comparable to those of the OVI animals (A1-A3). Stratum pyramidale (SP), Stratum oriens (SO) and Stratum radiatum (SR), Pyramidal cell layer (PO), Granular cell layer (G): Scale bars 50 μm.

#### 3.3.3. C1q expression

C1q immunoreactivity was observed in perikarya as well as dendritic processes of hippocampal neurons. In OVI animals, C1q +ve neurons of varied shapes and sizes were found mostly in SP of CA1 and CA3 (Fig. 5A1, A2) and granular and polymorphic cell layers of DG (Fig. 5A3). In OVX animals, on the whole, the staining intensity, as well as the density of C1q +ve neurons, was significantly higher in the various subfield of the hippocampus as compared to that of the OVI animals ((Fig. 5B1, B2, B3). E2 therapy to OVX rats resulted in significant downregulation of C1q in all the subfields of the hippocampus (Fig. 5C, C2; D1, D2), compared to that of the OVI like state (Fig. 5A1, A2). Downregulation of C1q irty was of similar intensity in CA1, and CA3 of TAM treated OVX animals as well (Fig. 5D1-D2). However, the downregulation of C1q irty in DG was somewhat less pronounced. Thus C1q +ve neurons appeared more intensely stained in DG of TAM treated group than in OVI and OVX+E2 treated groups.

**Fig. 5.**
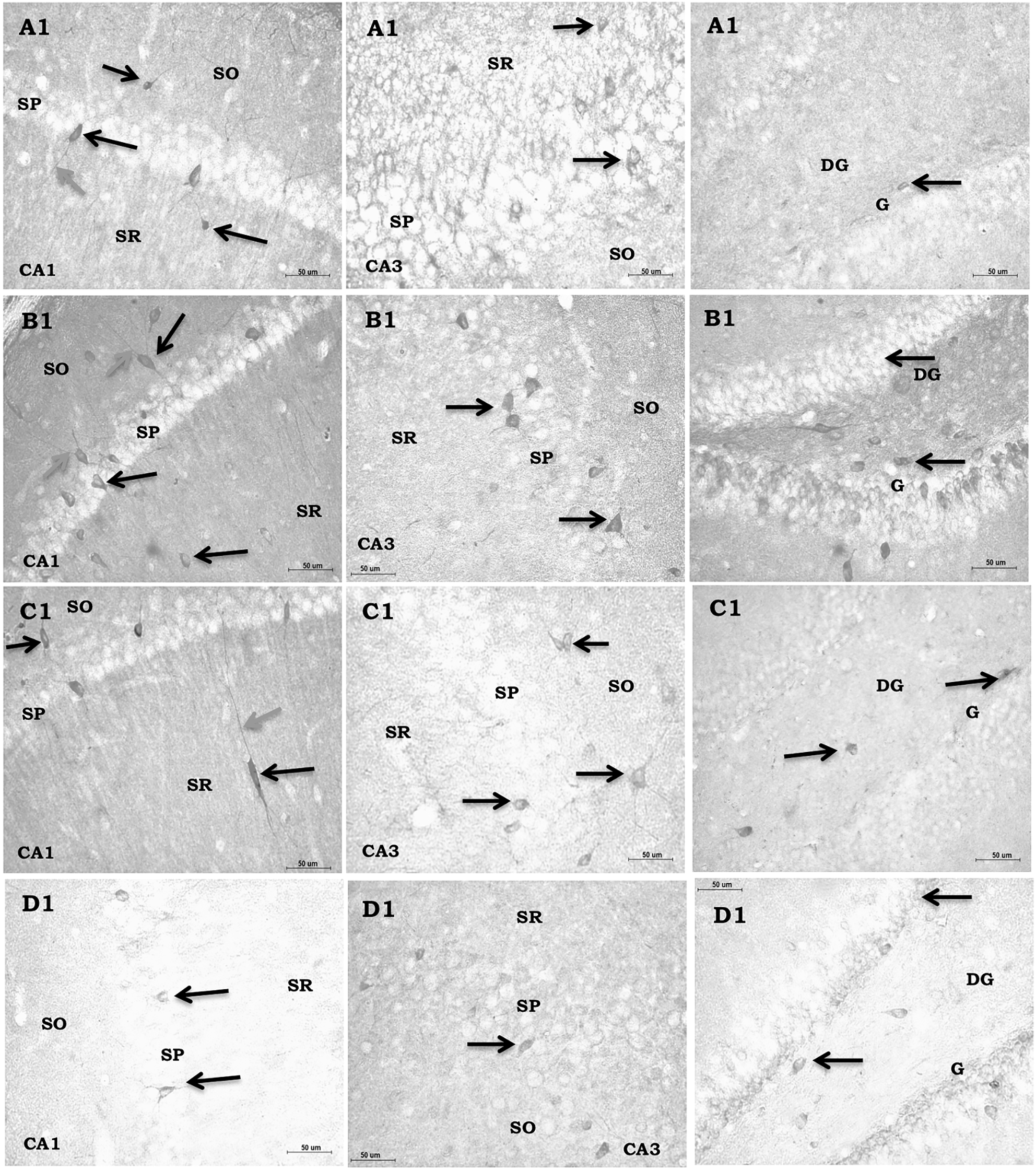
High-power photomicrographs of coronal sections of rat hippocampus (CA1, CA3 & DG) of OVI (A1-A3), OVX (B1-B3), OVX+ E2 (C1-C3) and OVX + TAM (D1-D3) showing distribution of complement C1q +ve neurons. Note: - Significant upregulation of C1q irty following ovariectomy (B1-B3). E2 (C) or TAM (D) therapy to OVX rats led to significant downregulation of C1q in all the hippocampal subfields, (C1-C3; D1-D3) the intensity being comparable to those of the OVI. Stratum pyramidale (SP), Stratum oriens (SO) and Stratum radiatum (SR). Pyramidal cell layer (PO) and Granular cell layer (G): Scale bars 50 μm.

#### 3.3.4. Quantitative analysis

IOD values for C3 [F (3, 16) = 10.52; P=0.0005; η2= 0.6636], C3aR [F (3, 16) = 6.278; P=0.005; η2= 0.5407], and C1q [F (3, 16) = 8.984; P=0.001; η2= 0.6275] in various hippocampal areas (CA1, CA3, and DG). OVX controls and OVX+Veh treated groups showed significantly higher values. Thereby suggestive of increased expression of these proteins in these groups as compared to ovary intact controls (Fig. 6A, B, C). IOD values for these proteins were significantly (p<0.01) lower in OVI, E2 treated, and TAM treated animals respectively (Fig 6A-C).

**Fig. 6.**
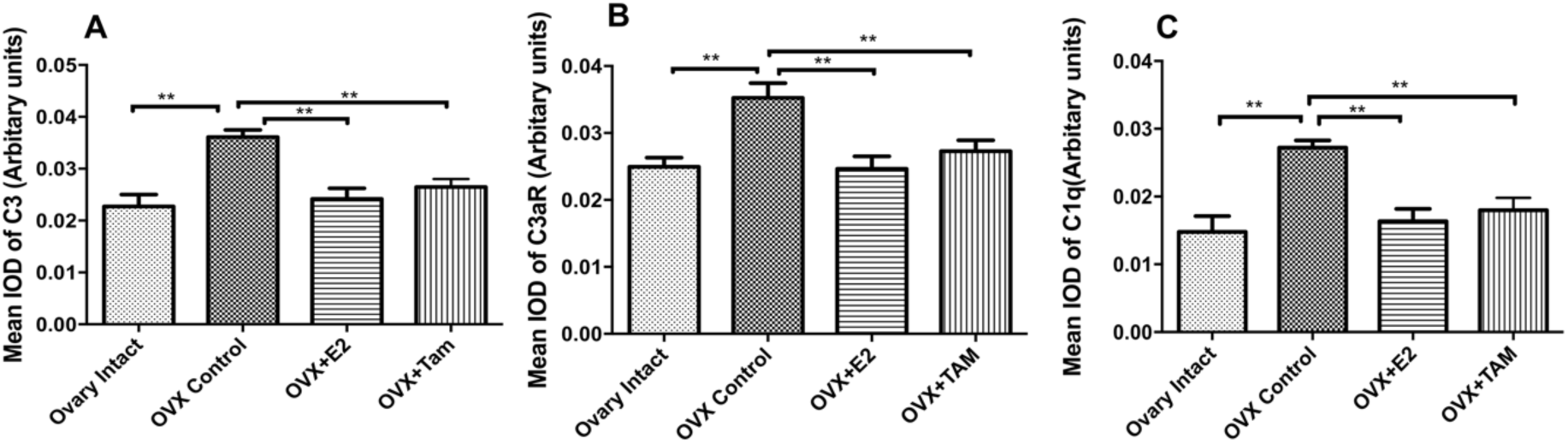
Histogram showing the mean IOD (Mean±SEM) of C3 (A), C3aR (B) and C1q-irty (C) in the hippocampus of experimental animals (**p<0.01), C3 and C1q-IOD was observed in OVX and OVX+Veh groups when compared to ovary intact controls., whereas IOD values in E2 and TAM treated OVX groups is comparable to ovary intact control levels.

### 3.4. Estrogen and tamoxifen therapy modulates mRNA, as well as protein levels

WB analysis was carried out to quantify C3 [F (4, 20) = 14.14; P<0.000; η2= 0.738], C3aR [F (4, 20) = 11.3; P<0.0001; η2= 0.693], and C1q [F (4, 20) = 9.904; P=0.001; η2= 0.664] proteins in the hippocampi of control and experimental animals. Immunoreactive bands for C3, C3aR, and C1q were visualized as ∼180kDa, ∼65kDa and ∼29Da protein respectively (Fig. 7D). A significant upregulation of C3, C3aR and C1q levels with OD (23.01±1.13; p = 0.0002), (25.62±1.01; p = 0.002) and (3.57±0.13; p = 0.002) respectively were observed in OVX animals as compared to ovary intact controls (14±1.25; 16.54±1.74 and 2.59±0.16). C3, C3aR, and C1q levels were significantly downregulated in E2 and TAM treated animals as compared to OVX controls, the OD being (16±1.21, p = 0.004; 16.59±1.68, p = 0.002; 2.60±0.17, p = 0.003) E2 treated group and (15.95±1.22, p = 0.003; 17.69±1.37; p = 0.008; 2.82±0.15 p = 0.03) in TAM treated group (Fig. 7B, C & D). However, the values were comparable between OV+Veh and OVX controls (23.29±0.92, p >0.999; 25.64±1.14, p >0.999; 3.55±0.15, p >0.999) (Fig 7B, C & D).

**Fig. 7.**
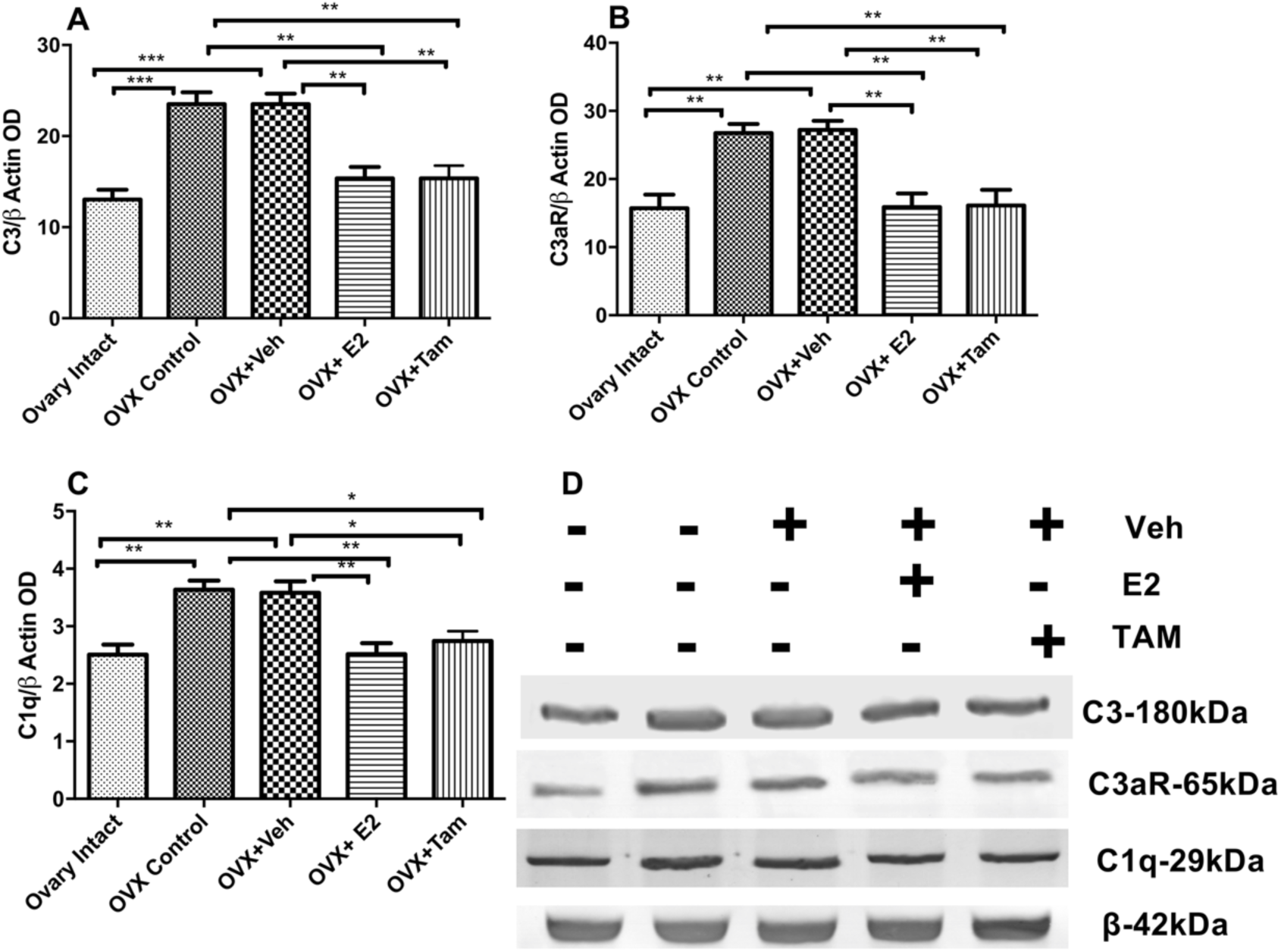
Histogram based on OD analysis of C3 (A), C3aR (B) and C1q (C) protein from the immunoblots (D) of hippocampus of control & experimental groups (n = 6). Densitometric analysis of C3, C3aR and C1q protein (**p<0.01; compared to OVX group). β-actin is used as an internal standard for equal loading of proteins.

In qRT PCR, GAPDH and β-Actin gene sequences were used as internal standards. Amplified gene products for C3 and C1q were detected in the hippocampi of all the control and experimental groups (Fig 8A, B & C).

**Fig. 8.**
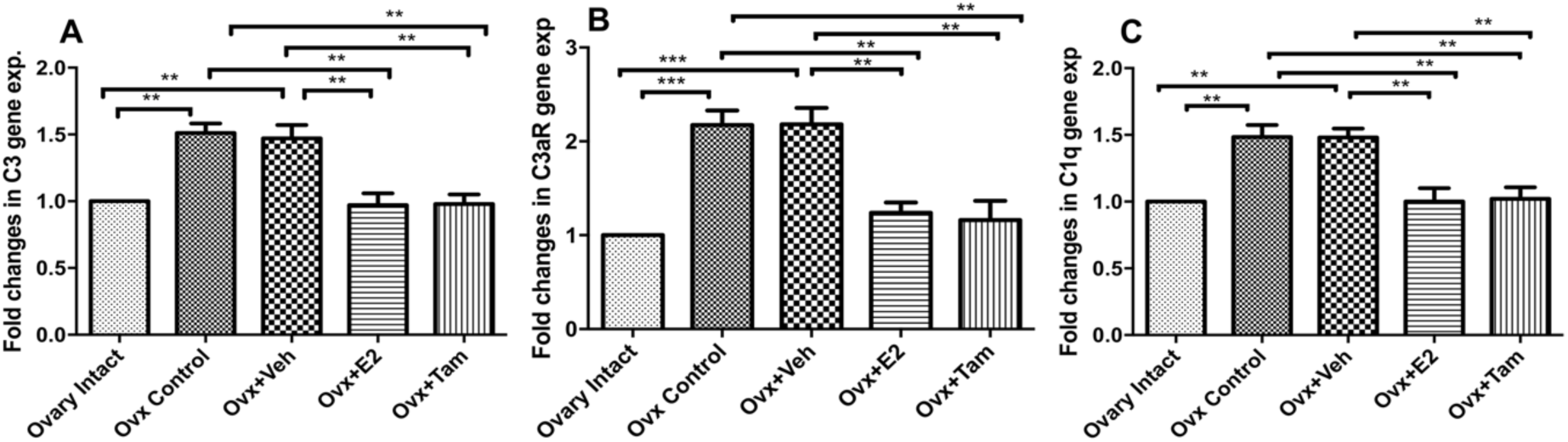
Histogram based on qRT-PCR expression analysis of C3 (A), C3aR (B) and C1q (C) showing the fold change of mRNA expression levels (C3, C3aR & C1q) in experimental samples relative to ovary intact control, while error bars show the standard deviation of ΔΔCT values obtained for the triple replicates of each sample type.

Hormone depletion (ovariectomy) led to significant upregulation of complement C3 [F (4, 20) = 11.24; P<0.0001; η2= 0.692], C3aR [F (4, 20) = 13.91; P<0.0001; η2= 0.735] and C1q [F (4, 20) = 11.24; P<0.0001; η2= 0.692] mRNA expression (Fig 8A, B & C) with a relative fold change (1.4±0.08, p = 0.002; 2.08±0.14, p = 0.008; 1.4±0.85, p = 0.002) respectively as compared to OVI controls. C3, C1q, and C3aR gene expression were significantly decreased following E2 and TAM therapy as compared to OVX controls, the relative gene expression of the above three complement proteins being (0.96±0.06, p = 0.001; 1.24±0.08, p = 0.003 and 1.0±0.07 p = 0.002) and (0.97±0.05p = 0.001; 1.27±0.19, p = 0.004 and 1.04±0.07, p = 0.006) respectively. C3, C3aR, and C1q mRNA levels were comparable between OVX+Veh and OVX controls the values being [(1.3±1.10 p >0.999; 2.08±0.16 p >0.999 and 1.43±0.06 p >0.999)] (Fig 8A, B & C).

### 3.5. Estrogen and tamoxifen therapy reduces neuronal loss

A significant increase in the TUNEL +ve cells was observed in all the subfields (CA1, CA3, and DG) of the hippocampus in OVX animals (Fig 9B). However, E2 and TAM therapy to OVX animals led to a decrease (∼9.17%; ∼11.5%) in the number of TUNEL +ve cells in all the subfields (CA1, CA3, and DG) (Fig 9D-E) as compared to OVX controls (19.7%) and ovary intact controls (∼5.33%, p<0.01) (Fig. 10).

**Fig. 9.**
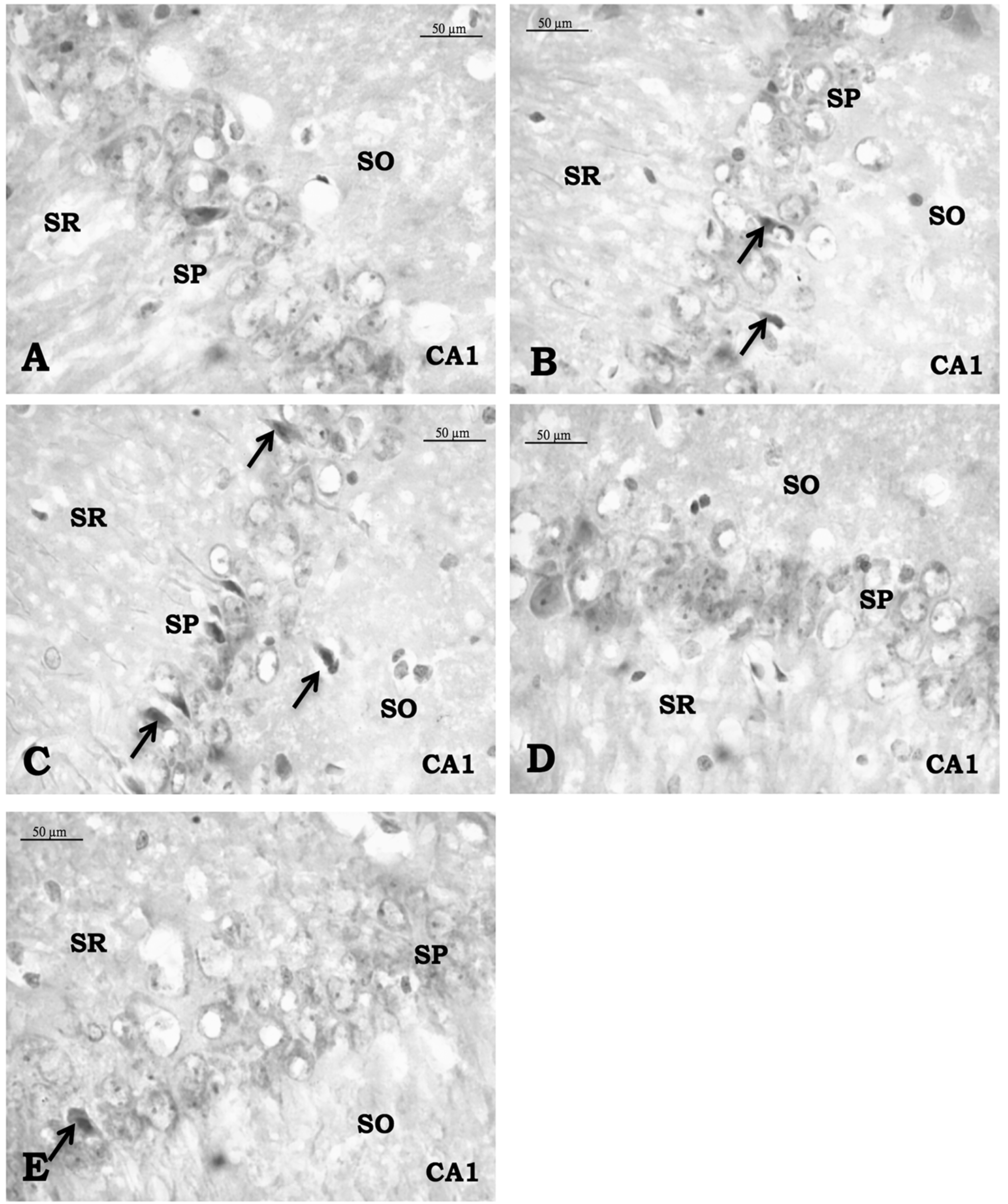
Representative high-power photomicrographs of coronal sections from ovary intact (A), OVX (B), OVX+Veh (C), OVX+E2 (D) and OVX+TAM (E) hippocampus showing the presence of TUNEL positive cells. For the identification of neuronal perikarya the sections were counterstained with a kit provided counterstain after TUNEL reaction. Arrows showing TUNEL positive apoptotic neurons. Scale bars 80μm. Stratum pyramidale (SP), stratum oriens (SO) and stratum radiatum (SR). Scale bars 50 μm.

**Fig. 10.**
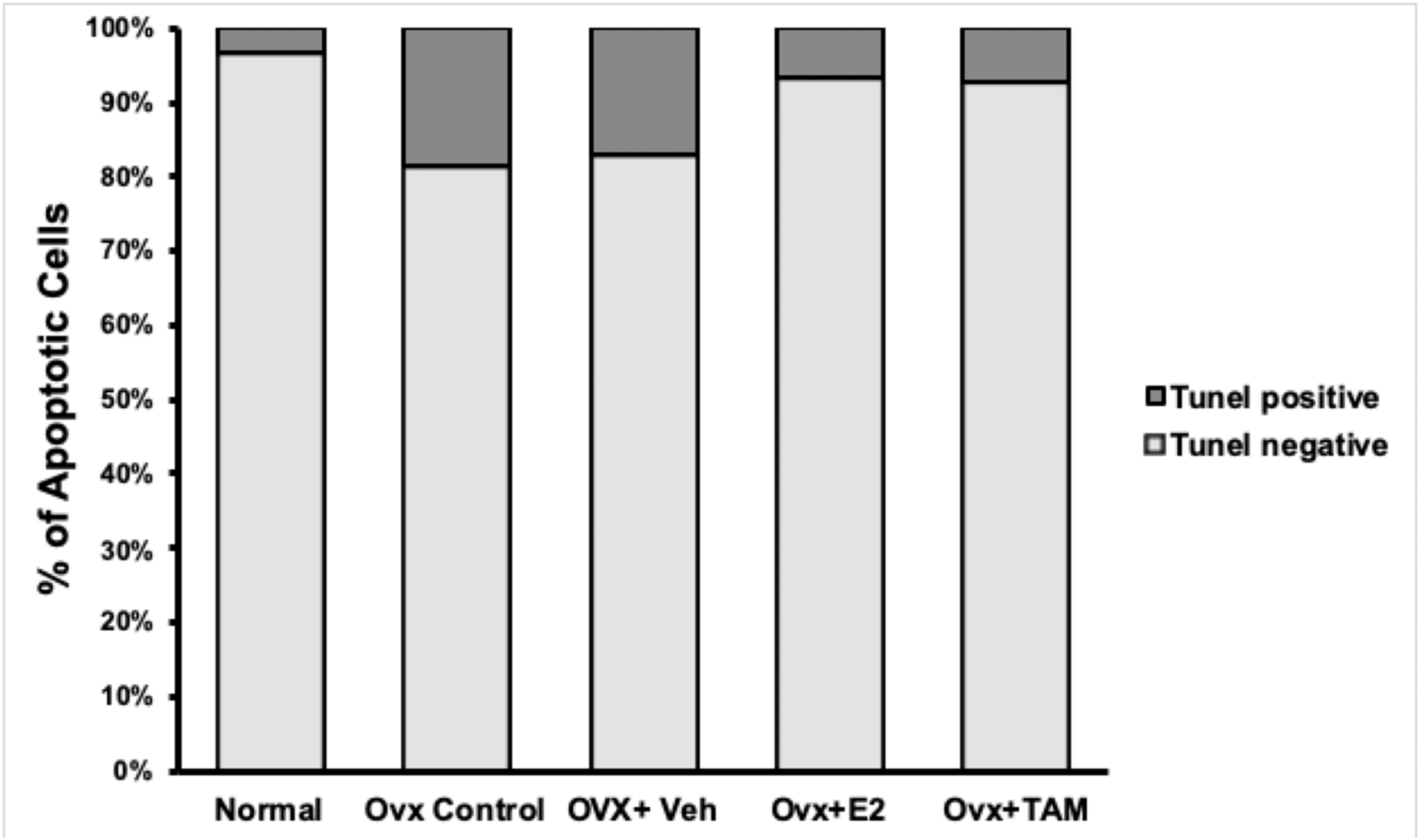
Histogram showing the percentage of apoptotic cells in the Hippocampus of various treated and control groups. (**p<0.01, relative to OVX control)

## 4. Discussion

Various cell types, including the neurons, have been reported to produce complement proteins. Using In situ hybridization and immunohistochemical techniques, a number of brain regions have been identified as the source of complement component and their receptors (Johnson et al., 1992; Shen et al., 1997; Singhrao et al., 1996; Spiegel et al., 1998; Terai et al., 1997). The role of the complement system in neurodegenerative diseases was initially suggested by Eikelenboom and co-workers (Eikelenboom et al., 1989). Since then, a good number of in vitro studies have consistently demonstrated the involvement of the complement pathway in the pathogenesis of neurodegenerative conditions such as AD (Akiyama et al., 2000). However, the role of the complement system in response to various insults in different areas of the brain including the hippocampus remains to be ascertained. Our observations regarding the influence of complement proteins at the genomic and proteomic level in the hippocampal area (memory and cognition) support the fact that the neurons are also the seat of essential complement protein expression. The present study provides preliminary evidence of the potential of estrogen and tamoxifen therapy in regulating the complement system levels (C1q, C3, and C3aR) in various sub-regions of the hippocampus of adult ovariectomized (OVX) female rats. Our observations are in line with previous studies reporting an increase in inflammatory events in reproductive as well as non-reproductive areas including the central nervous system following estrogen depletion in females (Vegeto et al., 2008). The perikaryal expression of C3 in CA1, CA3 and DG of ovariectomized animals (present study) was markedly high as compared to ovary intact controls. C3a is a polypeptide anaphylatoxin, expressed by tissue at sites of complement activation and works as a chemoattractant locally, C3aR mRNA expression in different brain regions has been reported in details by Davoust and co-workers (Davoust et al., 1999). Our present observation of C3aR receptor expression in pyramidal cell layer, SO and SR layer (CA1 and CA3) of OVI rat hippocampus are on the similar pattern as reported by Davoust et al (1999). In the present study, C3aR was predominantly expressed in the granular and polymorphic cell layer (DG) and similar pattern had been revealed by in situ hybridization studies carried out by Davoust et al (1999). Besides neurons, the hippocampal interneurons were also observed to be C3aR immunopositive. Upregulation of anaphylatoxin receptors in the brain cells has also been described in experimental models of focal cerebral ischemia (Barnum et al., 2002; Van Beek et al., 2000) and trauma (Stahel et al., 1997; Stahel et al., 2000), thereby suggestive of the role of C3aR in these pathologies.

Sarvari and co-workers (2011) demonstrated elevated expression of C1q in the frontal cortex of aged female rats, with low production of ovarian hormones. Sarvari et al. (2012) further reported on the abnormal proteins, such as amyloid-β and prion protein being recognized by C1q in the brain. The study carried out by Ricklin et al. (2010) demonstrated C1q binding to apoptotic cells, pointing towards initiation of the classical pathway of complement system into in phagocytosis of apoptotic cells and destruction of invading pathogens. In line with the study of Ricklin et al (2010) and Sarvari et al (2011), our observations also revealed upregulation of complement C1q in CA1, CA3 and DG of hippocampus in ovariectomized adult rats along with increased percentage of apoptotic cells, thereby indicative of increased inflammatory and neurodegenerative events induced by hormone depletion.

Activated microglia were also identifiable with the concomitant upregulation of a variety of cell-surface and cytoplasmic molecules like complement proteins, especially C1q. Our observations pertaining to the immunohistochemical expression of microglia to indicate their hyper ramification with following ovariectomy (**Supplementary Fig. S1 and S2**). However, E2 and TAM therapy restored glial morphology to the control status with low expression of C1q. Our observation do substantiate the reports of Raivich et al. (1999) and Block and Hong (2005) where in they have described the activated forms of microglia with characteristic morphological changes including the shortening of processes and a hypertrophied cell body.

Additionally, activated microglia do release several soluble pro-inflammatory factors like cytokines and complement components (Block and Hong, 2005; Colton and Gilbert, 1987; Liu et al., 2002; Sawada et al., 1989). Increasingly, chronic or deregulated inflammation is being recognized as a primary contributing factor to the progression and etiology of neurodegeneration (Barrientos et al., 2019; Bonifati and Kishore, 2007; Giovannini et al., 2002; Morale et al., 2006; Yun et al., 2018) as well as other disease conditions such as like atherosclerosis (Chait et al., 2005), arthritis (Mizuno, 2006) and osteoporosis (Clowes et al., 2005; Ginaldi et al., 2005). E2 has been demonstrated as one of the sex hormones that influences the progression of varied inflammatory pathologies including cardiovascular disease, (Hodgin and Maeda, 2002; Wagner et al., 2002), osteoporosis (Turner et al., 1994; Watts, 2002) and autoimmunity (Cutolo et al., 1995). Evidences from clinical trials and animal studies suggest the association of E2 in decreased incidence and delayed onset and progression of acute and chronic brain disorders ranging from stroke and schizophrenia to Alzheimer’s disease, multiple sclerosis, and Parkinson’s disease (Behl, 2002; Bisagno et al., 2003; Chowen et al., 2000; Lee and McEwen, 2001; Roof and Hall, 2000). The reduced susceptibility to acute brain injuries such as cerebral ischemia, neurotrauma and certain neurotoxic agents in pre-menopausal females, both in humans and rodents, strongly suggests a correlation between the levels of E2 and the severity and progression of inflammatory diseases.

Since hippocampus is considered as one of the first brain regions to be affected in AD with loss of synapses (perhaps the most important feature of the disease), investigations in the mechanisms of synapse elimination in the hippocampus are highly warranted, especially in females. In our current study on ovariectomized rats we have observed increase in mRNA levels and expression of inflammatory markers along with concomitant decrease in the synaptophysin (presynaptic marker) mRNA and protein in the hippocampus following estrogen depletion (**Supplementary Fig. S3 and S4**). These results are in good agreement with the earlier observation of Stevens et al. (2007) and Chu et al. (2010) who reported involvement of complement system in the elimination of retinogeniculate and sensorimotor cortex synapses. A role for microglia has been suggested based on observation inhibited microglial motility inducing defective synapse elimination in the hippocampus (Paolicelli et al., 2011). In addition, Paolicelli and colleagues (2011) have also reported the presence of postsynaptic density protein PSD95 in microglia, suggestive of synaptic protein being engulfed by these cells.

The hormone replenishment therapy has been suggested to induce an anti-inflammatory reaction (Abu-Taha et al., 2009; Vegeto et al., 2008). Estrogen has been reported to play a modulatory role in animal models of various neuroinflammatory diseases (Brown et al., 2010; Lewis et al., 2008; Mor et al., 1999; Sarvari et al., 2012; Tiwari-Woodruff et al., 2007; Vegeto et al., 2003; Yun et al., 2018). Our study aimed at determining the effect of ovariectomy & hormone replenishment on the expression of complement proteins and the observations revealed a significant difference in the complement expression in OVX and treated groups (E2 or SERM), the expression level in the later being comparable to the OVI group. Perikaryal expression of C3 in the pyramidal cell layer of CA1 and CA3 and polymorphic and granular layers of DG was significantly downregulated following hormone replenishment in OVX animals. The observations pertaining to C1q expression also presented a similar pattern, with upregulated expression of C1q in OVX animals therapy bringing about downregulated expression of C1q following E2 replenishment, the expression being comparable to that of OVI level.

Absence of C1q accompanied by a significant reduction of inflammatory reaction in the CA3 region of the hippocampus in a transgenic rat model was reported by Fonseca et al (2004) and McGeer & McGeer (1998). In our ovariectomized rat model, a significant decrease in C1q expression was observed in both the SP as well as in SO layer of CA1 and CA3 following E2 therapy. These observations do support the modulatory role of estrogen on neurodegenerative and inflammatory events in the CNS. This presumption was further confirmed with TUNEL assay which showed an increase and a decrease in the number of apoptotic cells following hormone depletion and E2 therapy respectively, thereby suggestive of neuroprotective effect of E2 (Sharma and Mehra, 2008).

Further validation of IHC expression of above mentioned markers with Western blot and Real-time quantitative PCR (qRT-PCR) analysis revealed increased C3 and C1q gene expression in the hippocampus of OVX animals. Interestingly, earlier reports have documented 1.8 and 4.3-fold increase of C1q and C3 mRNA levels in the frontal cortex of middle-aged rats but not in young adult rats following ovariectomy (Sarvari et al., 2012). C1q mRNA has been reported to get increased by 11-80 fold over control levels, indicating complement component production in the affected areas (entorhinal cortex, hippocampus, and mid-temporal gyrus) of the AD brain (Yasojima et al., 1999). During neuronal inflammation, increase in complement proteins and mRNA levels in resident brain cells in different inflammatory disease animal models has also been reported (Dietzschold et al., 1995; Stahel et al., 1998; Stahel et al., 1997).

Tamoxifen is a synthetic non-steroidal compound and because of its ability to cross the blood-brain barrier, the effects of tamoxifen on inflammatory protein and mitochondrial function have been reported in various brain pathologies eg, cerebral ischemia, subarachnoid hemorrhage, and spinal cord injury (Baez-Jurado et al., 2019; Franco Rodriguez et al., 2013; Moreira et al., 2004, 2005). Tamoxifen administration stimulates cholinergic function, thereby improving learning and memory in postmenopausal women. During the Virtual Morris Water Maze (VMWM) task enhanced cholinergic system function and modulation of hippocampal activity was reported in ovariectomized rats (OVX) by protection of oxidative damage of the brain (Newhouse et al., 2013; Zabihi et al., 2014). Reduction of reactive astrocytes could be mediated by protein kinase inhibition; whereas, decreased levels of cytokines, mediated by ERα and NF-κB could be promoting neuronal survival in this type of injury (Horgan et al., 1986). Tamoxifen administration to TBI rat reduced apoptosis and improved functional outcomes by activation of the p-ERK1/2 signaling pathway and BCl2 expression (Tsai et al., 2014). Tamoxifen may bind to the GPR30 receptor and activate the kinase signaling pathway (Akt, ERK, and PI3K), in turn being helpful in improving neuronal health (Arevalo et al., 2015; Tang et al., 2014).

Estrogen and tamoxifen-activated estrogen receptors have been reported to stimulate the promoter region of the complement C3 gene (Fan et al., 1996). The observations of the present investigation also showed concomitant upregulation of both C3 and C1q in ovariectomized rat hippocampus, and their reversal near to OVI like state following E2 & TAM therapy is also in line with these suggestions.

A very suitable rationale conclusion of the present investigations is that estrogen depletion leads to neurodegenerative and neuroinflammatory events in female rat brain, and these increased inflammatory events further exaggerate the neurodegeneration via complement proteins C1q and C3.

## Acknowledgements

This work was partially supported by ICMR Fellowship 45/13/2011/IMM/BMS (Senior research fellow) and DST N-PDF Fellowship (National Postdoctoral fellowship), Delhi, India.

---

**Fig. S1.**
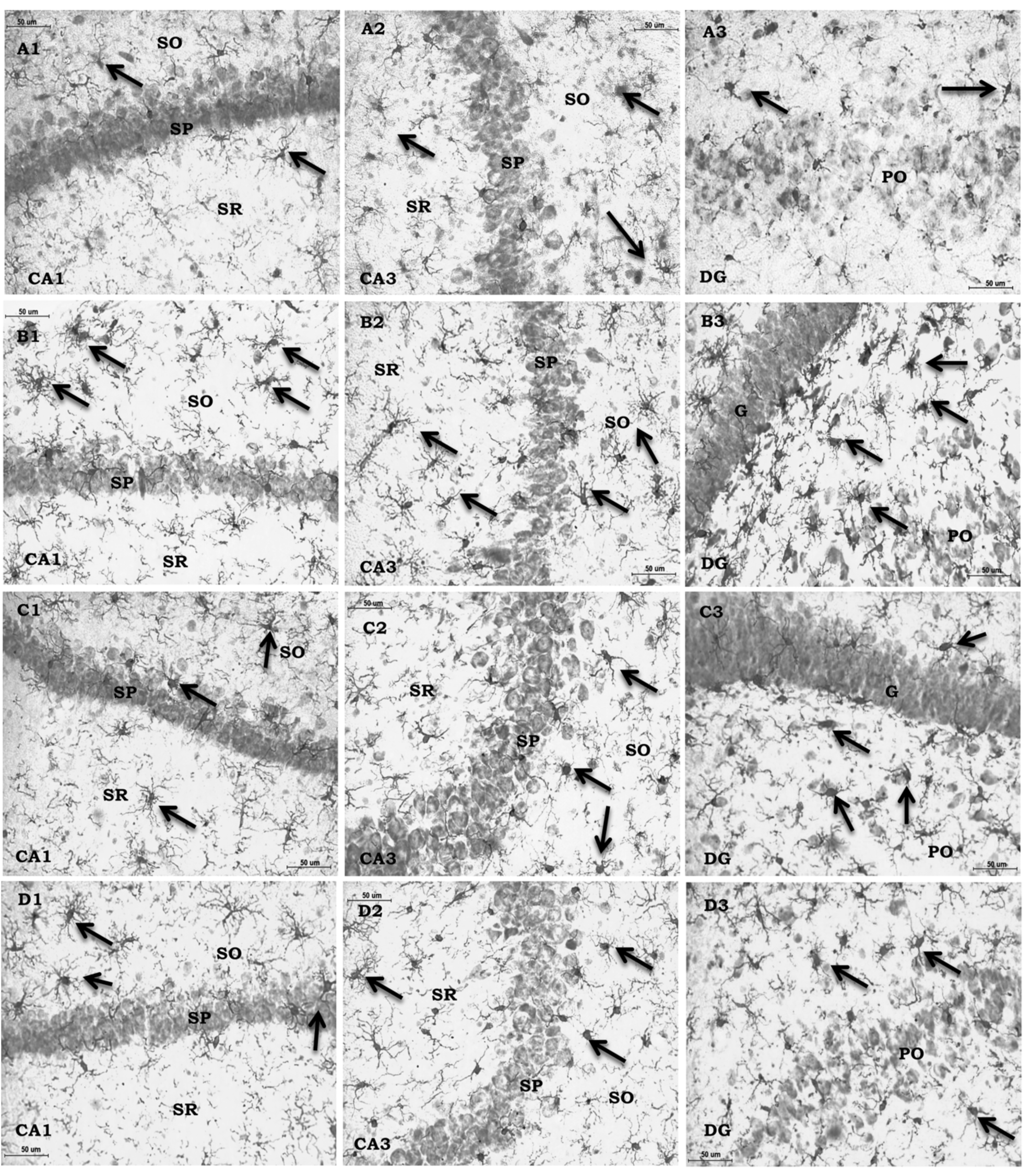
High-power photomicrographs of coronal sections of rat hippocampus (CA1, CA3 & DG) of OVI (A1-A3), OVX (B1-B3), OVX+ E2 (C1-C3) and OVX + TAM (D1-D3) showing expression of Iba1. Note: - Following ovariectomy the Iba-1immunoreactivity (irty) was significantly upregulated (B1-B3). E2 (C) or TAM (D) therapy to OVX rats led to significant downregulation of Iba1 in all the hippocampal subfields, (C1-C3; D1-D3) the intensity being comparable to those of the OVI. Stratum pyramidale (SP), Stratum oriens (SO) and Stratum radiatum (SR). Pyramidal cell layer (PO) and Granular cell layer (G): Scale bars 50 μm.

**Fig. S2.**
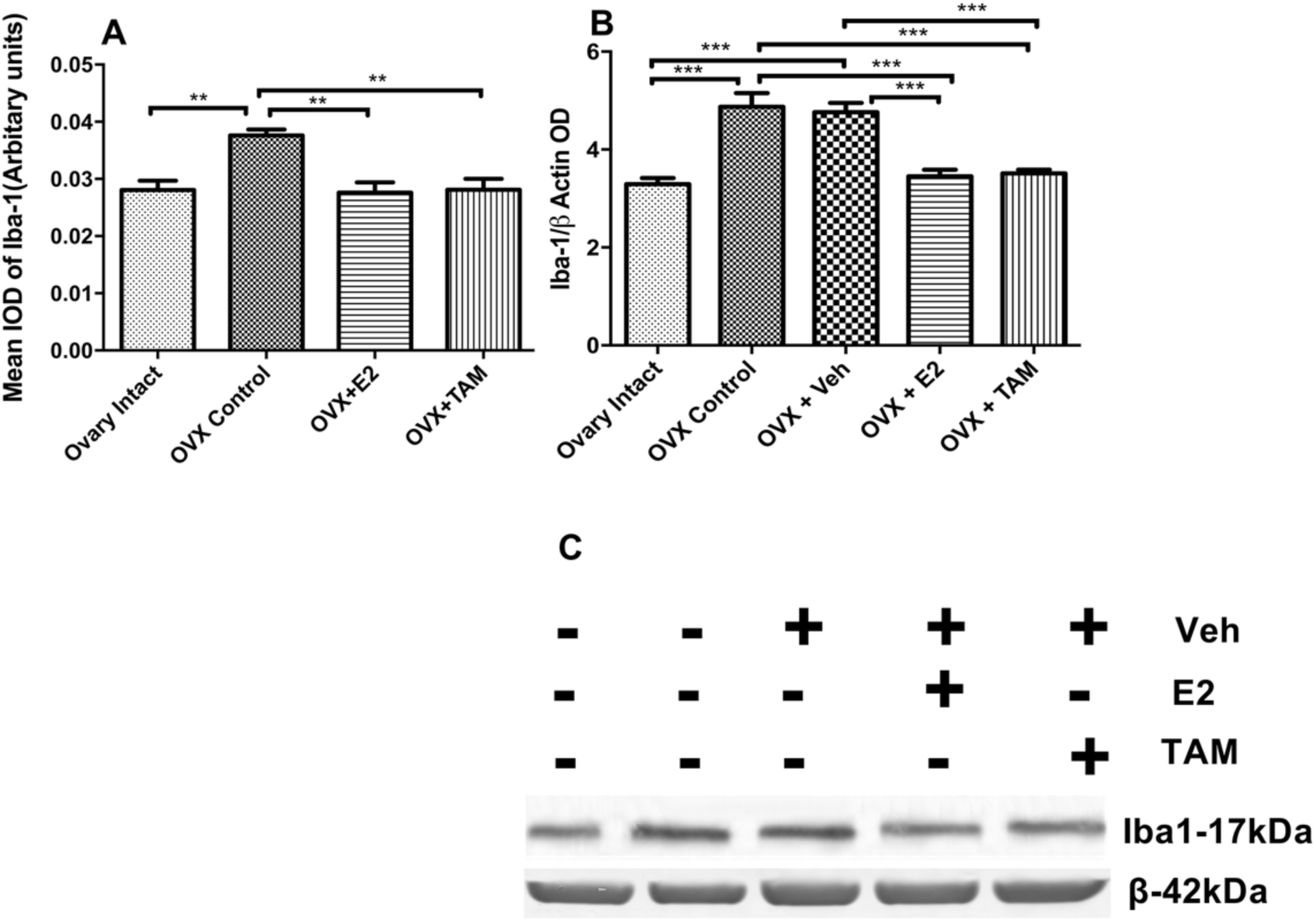
Histogram showing the mean IOD (A) and WB OD (B&C) levels of microglia (Iba1) (Mean±SEM) in the hippocampus of experimental animals (**p<0.01 and ***p<0.001), Iba1 was observed in OVX and OVX+Veh groups when compared to ovary intact controls, whereas values in E2 and TAM treated OVX groups is comparable to ovary intact control levels.

**Figs. S3.**
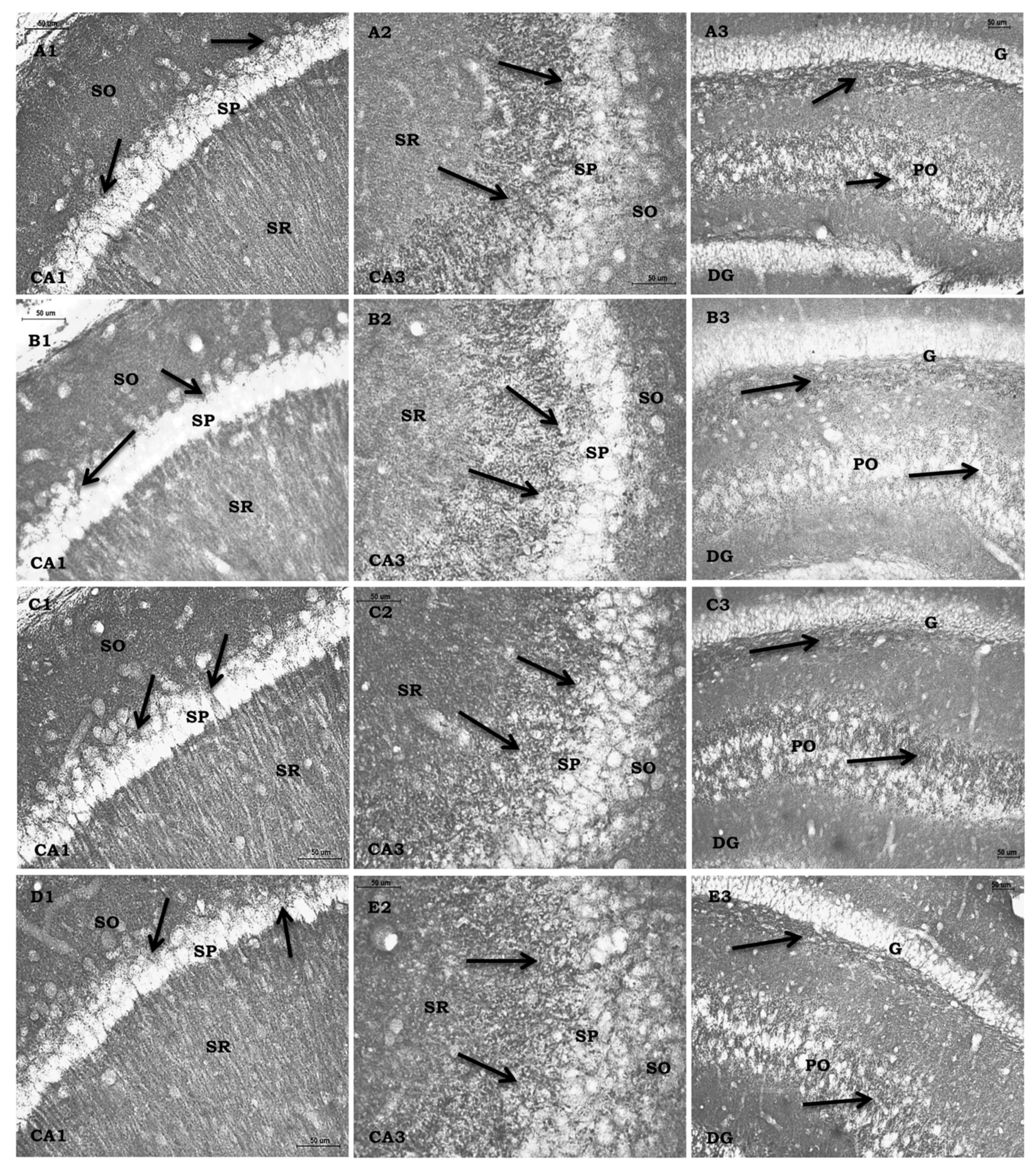
Representative high-power photomicrographs of coronal sections of rat hippocampus (CA1, CA3 & DG) of OVI (A1-A3), OVX (B1-B3), OVX+ E2 (C1-C3) and OVX + TAM (D1-D3) showing distribution of SYP-irty. The arrows point to bouton-like structures in all the subfields of the hippocampus. Note: Following ovariectomy the SYP-irty was significantly down-regulated (B1-B3). E2 (C) or TAM (D) therapy to OVX rats led to significant upregulation of SYP in all the hippocampal subfields, (C1-C3; D1-D3) the intensity being comparable to those of the OVI. Stratum pyramidale (SP) and stratum oriens (SO). Stratum radiatum Scale bars 50 μm

**Fig. S4.**
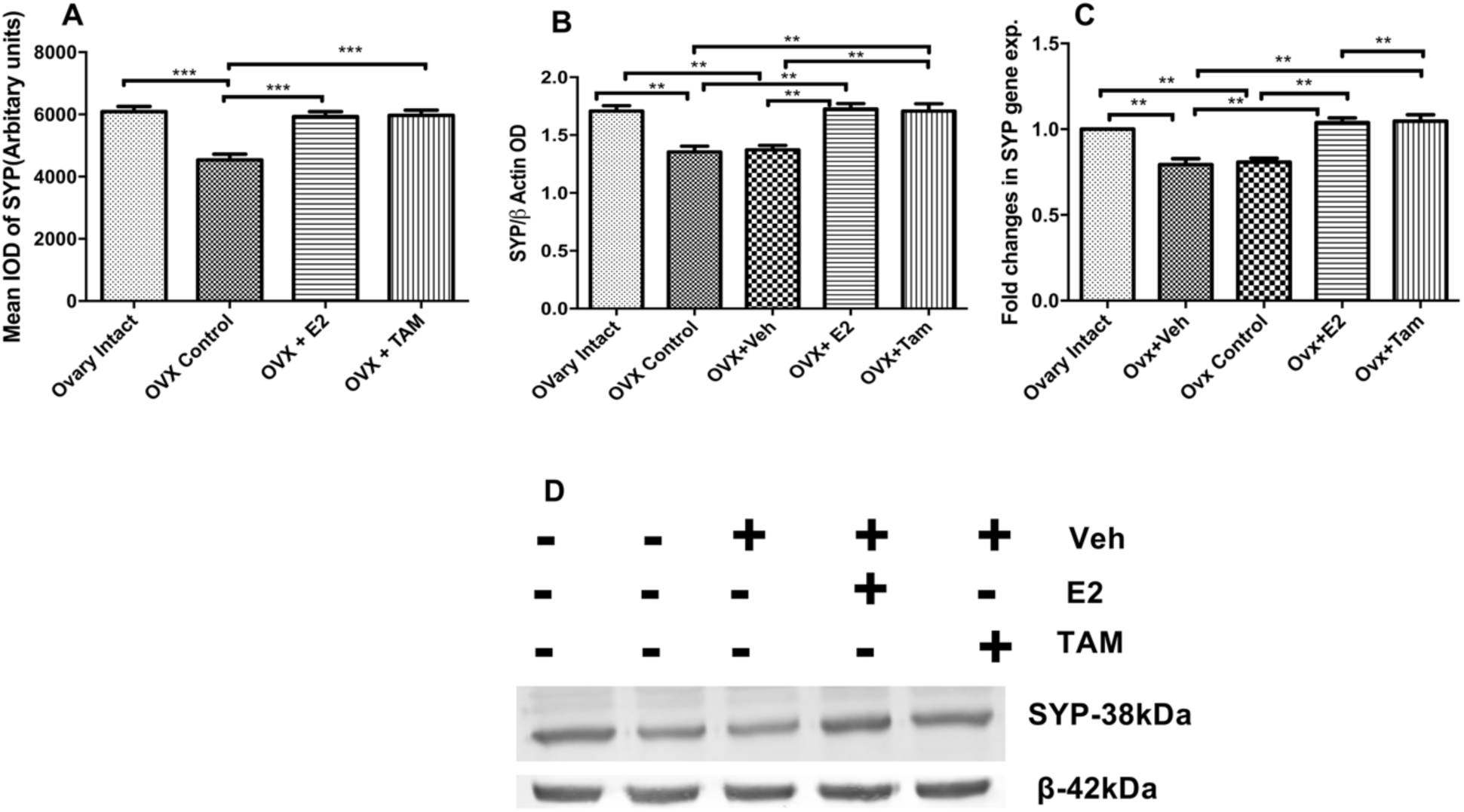
Histogram showing the mean IOD (A), WB OD (B&D) and mRNA (C) levels of synaptophysin (SYP) (Mean±SEM) in the hippocampus of experimental animals (**p<0.01 and ***p<0.001), SYP was observed in OVX and OVX+Veh groups when compared to ovary intact controls, whereas values in E2 and TAM treated OVX groups is comparable to ovary intact control levels.

